# Context-aware sequence-to-function model of human gene regulation

**DOI:** 10.1101/2025.06.25.661447

**Authors:** Ekin Deniz Aksu, Martin Vingron

## Abstract

Sequence-to-function models have been very successful in predicting gene expression, chromatin accessibility, and epigenetic marks from DNA sequences alone. However, current state-of-the-art models have a fundamental limitation: they cannot extrapolate beyond the cell types and conditions included in their training dataset. Here, we introduce a new approach that is designed to overcome this limitation: Corgi, a new context-aware sequence-to-function model that accurately predicts genome-wide gene expression and epigenetic signals, even in previously unseen cell types. We designed an architecture that strives to emulate the cell: Corgi integrates DNA sequence and *trans*-regulator expression to predict the coverage of multiple assays including chromatin accessibility, histone modifications, and gene expression. We define *trans-*regulators as transcription factors, histone modifiers, transcriptional coactivators, and RNA binding proteins, which directly modulate chromatin states, gene expression, and mRNA decay. Trained on a diverse set of bulk and single cell human datasets, Corgi has robust predictive performance, approaching experimental-level accuracy in gene expression predictions in previously unseen cell types, while also setting a new state-of-the-art level for joint cross-sequence and cross-cell type epigenetic track prediction. Corgi can be used in practice to impute context-specific assays such as DNA accessibility and histone ChIP-seq, using only RNA-seq data.

## Main Text

A central goal of genomics is to understand how the genome dictates its functional outputs, such as gene expression and epigenetic state. To this end, sequence-to-function models have been highly successful in making these predictions from DNA sequences alone^1^. Trained on the human and mouse reference genomes and large functional genomics datasets, these models learn gene regulatory grammar and can accurately predict the activity of previously unseen sequences^2–4^. They are useful in predicting effects of non-coding variants and generating synthetic sequences^5–7^. However, current state-of-the-art models have a fundamental limitation: they cannot extrapolate beyond the cell types and conditions included in their training dataset. Since they operate on DNA sequence alone, these models are blind to the biological context in which the sequence is interpreted.

Context-aware sequence-to-function models overcome this fundamental limitation by enabling generalization to previously unseen cellular contexts. These models operate on not only the DNA sequence but also a context vector. The context vector is computed from knowledge about the cell state: usually gene expression or chromatin accessibility. Previous models used transcription factor (TF) expression^8–11^, ATAC-seq^12,13^ and single-cell multiome (joint RNA and ATAC)^14^ data. However, these models are limited either in their predictive performance, low number of predicted assay types, or reliance upon known TF binding motifs.

Here we introduce **Corgi**, a new context-aware sequence-to-function model that predicts genome-wide coverage of 16 different assays including RNA-seq, chromatin accessibility, DNA methylation and histone modifications in human cells. Corgi stands for “**Co**ntext-aware **R**egulatory **G**enomics **I**nference” and it continues the canine-themed naming tradition in sequence-to-function models. Importantly, we designed Corgi’s architecture to imitate cellular gene regulation. In the cell, *trans-*regulatory proteins bind DNA and RNA to drive transcription, modulate chromatin state and control mRNA decay. So for its context vector, Corgi uses the expression of *trans-*regulators, which are defined as transcription factors, transcriptional co-activators, chromatin modifiers, and RNA binding proteins.

We address a central challenge in context-aware sequence-to-function models: it is not straightforward to integrate DNA sequence with the context vector due to the difference in their dimensions. The context vector is one-dimensional, while the DNA sequence has two dimensions: length and features. We leverage the feature-wise linear modulation^15^ (FiLM) technique to combine the two. FiLM was previously employed in various tasks such as image generation^16^ and speech recognition^17^, and has proven technically and conceptually appropriate for this task.

After training on a diverse set of human bulk and single-cell contexts, we show that Corgi can accurately generalize to previously unseen contexts. Corgi approaches experimental-level accuracy in predicting gene expression in new cell types, while accurately predicting DNase-seq, ATAC-seq, DNA methylation and various histone marks. Remarkably, Corgi’s performance remains robust even on the toughest benchmarks which test the model on held-out sequences and held-out cell types, where Corgi sets a new state-of-the-art prediction accuracy. Furthermore, we show that Corgi has increased imputation accuracy compared to tensor decomposition-based methods when predicting epigenetic tracks from RNA-seq, and has learned aspects of cell type-specific gene regulation.

## Results

### Model architecture and training data

Corgi builds upon a hybrid convolutional-transformer architecture, similar to recently published DNA sequence models^18^. The input 524 kb one-hot encoded DNA sequence is first processed through several layers of convolution and max pooling operations, aiming to extract meaningful sequence features and to reduce the length of the sequence by decreasing resolution (Fig. 1). This step computes the *cis*-features matrix that summarizes the local *cis*-regulatory landscape. To enable context-awareness, Corgi has a second input: the *trans*-regulatory context vector. This vector represents expression levels of selected *trans-*regulator genes, and is processed using a multilayer perceptron which computes the *trans*-features. Subsequently, *cis*- and *trans*-features are integrated by the FiLM layers, which apply an affine transformation to the *cis*-features matrix along its feature axis. These are followed by transformer layers that aim to capture long-range genomic interactions such as enhancer-promoter loops through self-attention. The final layer outputs coverage predictions for 16 different genomic assays at a 64 bp resolution. For a more detailed description of the architecture, see Methods and Figure S1.

**Figure 1.**
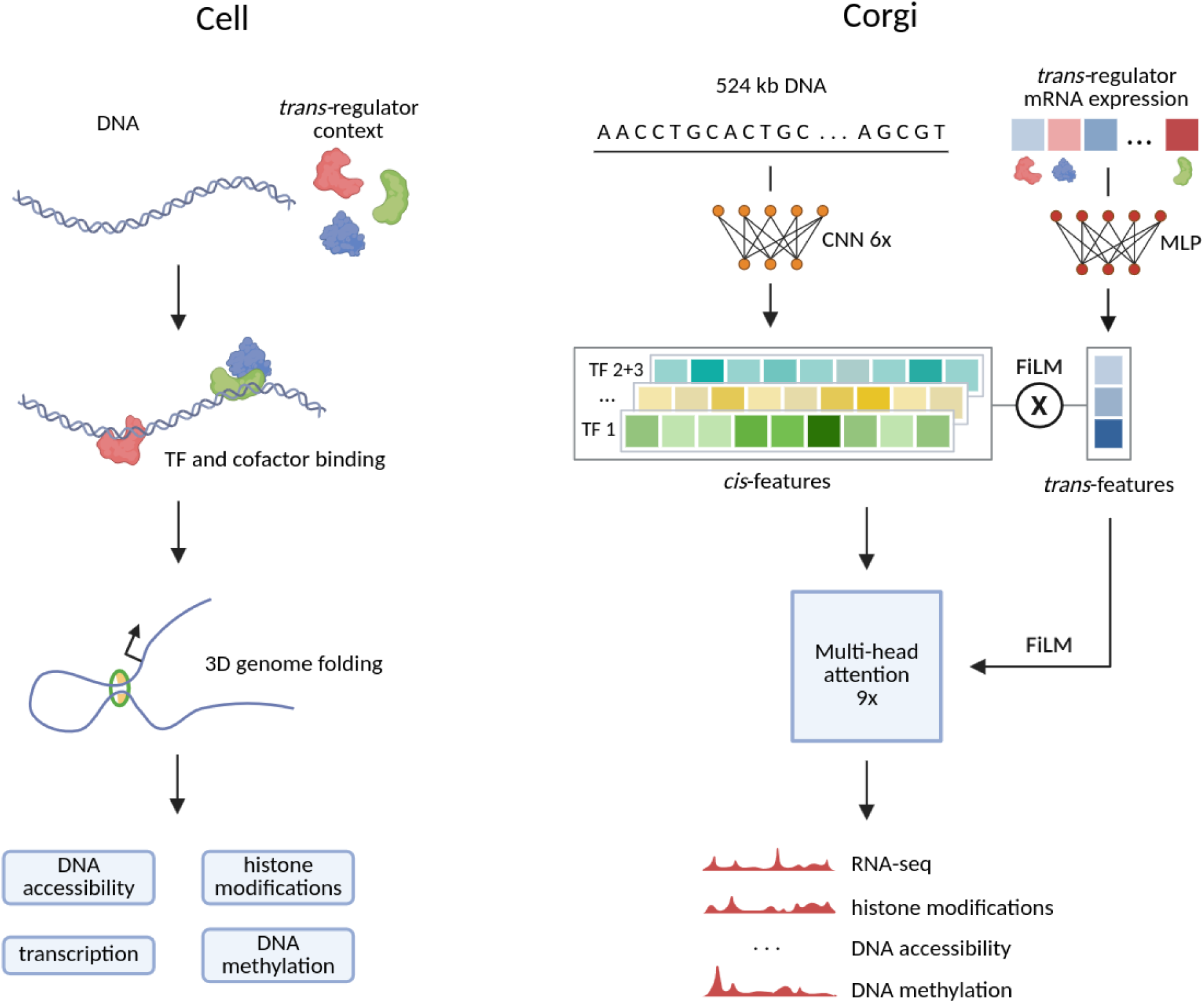
Corgi’s architecture strives to imitate cellular gene regulation. We designed an architecture that strives to emulate how cells read the regulatory genome using *trans* factors. The model first extracts *cis*-regulatory features via convolutional layers and *trans*-regulatory features via an MLP operating on *trans-*regulator expression. These are integrated using FiLM layers, which apply feature-wise affine transformations akin to biophysical “affinity × concentration” models. Instead of single TFs, these features are combinations of *trans-*regulators learned by the model. Afterwards, transformer blocks with FiLM layers capture long-range genomic interactions in a cell-type specific manner.

This design mirrors cellular gene regulation: for a regulatory element to be active, both a *cis*-element and its corresponding *trans*-regulator must be present. Mathematically it is a multiplication of *cis*- and *trans*- features, and if either one of them is zero, the product is also zero. This is conceptually similar to models like activity-by-contact (ABC)^19^, which multiply activity with contact. In Corgi, these features can be combinations of different factors that are learned during the training process. There are FiLM layers within the convolutional blocks as well as the transformer blocks, ensuring that *trans-*regulatory context can be integrated into the sequence at multiple points.

We trained Corgi on a large dataset of sequencing experiments from ENCODE^20^, FANTOM5^21^, Tabula Sapiens^22^, CATlas^23^ after careful curation (Methods). The dataset includes DNase-seq, ATAC-seq, RNA-seq, ChIP-seq and whole-genome bisulfite sequencing (WGBS) experiments. We used a very strict training-test split, to minimize potential data leakage between training, validation and test contexts (Figure S2, Methods).

### Corgi accurately predicts genomic tracks in held-out sequences and contexts

In order to use the diverse training set most efficiently, we first harmonized input *trans-*regulator expression levels that were obtained by different transcriptomic assays by quantile normalization, log transformation and batch correction. We assessed the impact of harmonization by focusing on samples with at least two different transcriptomic assays available. Harmonization reduced batch effects between different assay types, and improved correlation between assays with the most striking examples being CAGE-seq vs RNA-seq (Figure S3).

Then, we evaluated Corgi’s performance under three different benchmarking regimes: cross-cell type, cross-sequence, and cross-both. In the cross-cell type setting, the model predicts on held-out cell types using sequences seen during training, thus testing generalization to new biological contexts. In the cross-sequence benchmark, predictions are made on unseen genomic regions within training cell types. This is the only benchmark that traditional non-context-aware models can be evaluated in. Finally, the stringent cross-both setting evaluates predictions on both unseen cell types and unseen sequences. This benchmark represents a very hard challenge for sequence-to-function models, as accurate prediction requires generalization capabilities across new cell types and new sequences. For all benchmarks we applied test-time data augmentation by averaging predictions for slightly shifted and reverse complemented sequences.

Corgi has strong overall performance across a multitude of experiments (Figure 2). In the cross-cell type setting, Corgi achieves an average Pearson’s *r* of 0.84 for DNase-seq predictions across genomic bins. The performance changes considerably across assays: histone modification ChIP-seq tracks show the highest variability ranging from 0.39 for H3K9me3 up to 0.83 for H3K4me3 (Figure S5). The model also predicts gene expression very accurately, achieving an average of 0.59 for CAGE and 0.79 for bulk RNA-seq tracks. Single-cell RNA-seq cannot be predicted as well as the bulk experiments, probably due to low abundance of data in our training set and possibly due to its inherent sparsity. Finally, DNA methylation can be predicted with very high accuracy (mean *r* 0.92), and in some samples approaching near-perfect predictions.

**Figure 2.**
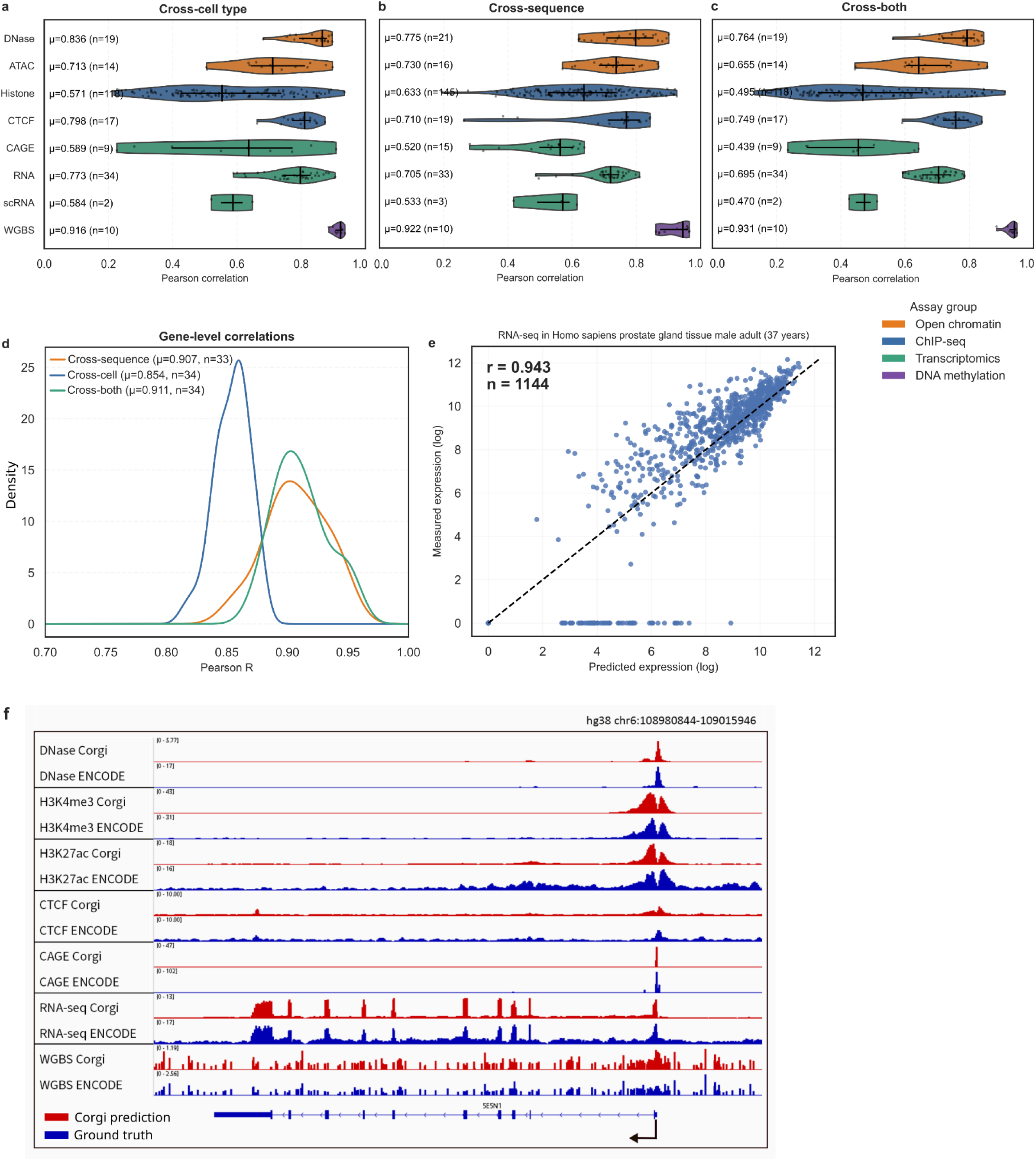
Genomic coverage prediction performance across different tracks. **(a), (b) and (c)** Violin plots showing model performance across different assays in cross-cell type, cross-sequence and cross-both settings. Correlations between predictions and ground truth data across genomic bins are reported, each data point is a correlation coefficient in one assay. **(d)** Density plots showing the performance distribution in gene-level RNA-seq predictions. Values in exon-overlapping bins were summed and log-transformed to calculate gene-level scores. **(e)** Scatterplot showing the relationship between predicted and measured log expression in the prostate tissue in genes from the test set. Each point represents the expression level of one gene, and the dashed line represents perfect predictions. **(f)** Corgi predictions (red) and ENCODE ground truth data (blue) are shown in an example locus from chr6 containing the SESN1 gene in sample #323 (testis), from the cross-both benchmark. Remarkably, the model captures histone mark signal shapes, the site of the CAGE-signal, DNA methylation patterns, and exact exon-intron boundaries in RNA-seq.

In the cross-sequence setting, Corgi maintains strong predictive performance, with increases in performance in 10 assays including all histone modifications, and modest drops in performance in seven assays. DNA accessibility and bulk RNA-seq predictions are not far from their cross-cell type counterparts, with 0.78 and 0.71 respectively.

Under the most challenging cross-both setting, which involves generalization to both unseen sequences and unseen cell types, Corgi still performs robustly, reaching 0.76 for DNase and 0.69 for RNA-seq. DNA methylation is predicted again with very high accuracy, reaching 0.93.

For gene expression prediction, it is often more informative to use log-transformed TPM values instead of the exact RNA-seq coverage across the gene. To simulate this, we computed gene-level expression predictions by aggregating coverage predictions over annotated exons and applied a log-transformation. Corgi achieves an average gene-level Pearson’s *r* value of 0.85 in the cross-cell type setting, and 0.91 in the cross-sequence and cross-both settings (Figure 2d). Importantly, Corgi’s performance is robust in different benchmarks and does not drop off when generalizing to held-out sequences or cell types.

A representative genome browser view illustrates Corgi’s predictive capacity across a 35 kb region on chromosome 6, encompassing the *SESN1* locus (Figure 2f). This example is taken from the cross-both setting, which shows a held-out sequence in a held-out cell type. The model faithfully reconstructs a wide range of assays, including chromatin accessibility, histone modifications, transcription initiation, gene expression, and DNA methylation. Corgi captures both the sharper peaks characteristic of DNase hypersensitive sites as well as the broader domains marked by active histone modifications such as H3K4me3 and H3K27ac. The predicted expression landscape across *SESN1* closely mirrors experimental RNA-seq and CAGE signals, with the exon-intron boundaries accurately predicted in the RNA-seq track and the exact TSS predicted in the CAGE track. Finally, the model can also recapitulate DNA methylation patterns with high precision.

### Corgi improves the state-of-the-art in context-aware epigenetic predictions

We benchmarked Corgi against EpiGePT, a state-of-the-art context-aware model for epigenetic track predictions. EpiGePT is the latest in a line of models that use DNA sequence and TF expression to predict epigenetic tracks. It uses the RNA expression values of transcription factors as its cell state vector, similar to Corgi. However, EpiGePT also relies upon known TF binding motifs to scan the input DNA sequence and calculate binding affinities.

Corgi clearly surpassed EpiGePT in both Pearson and Spearman correlation metrics (Figure 3a, b), reaching an average of 0.38 versus 0.08 on histone modification ChIP-seq tracks. Furthermore, visual inspection in the genome browser shows that EpiGePT predictions have a higher noise and DNase-seq peaks are often not correctly identified (Figure 3e).

**Figure 3.**
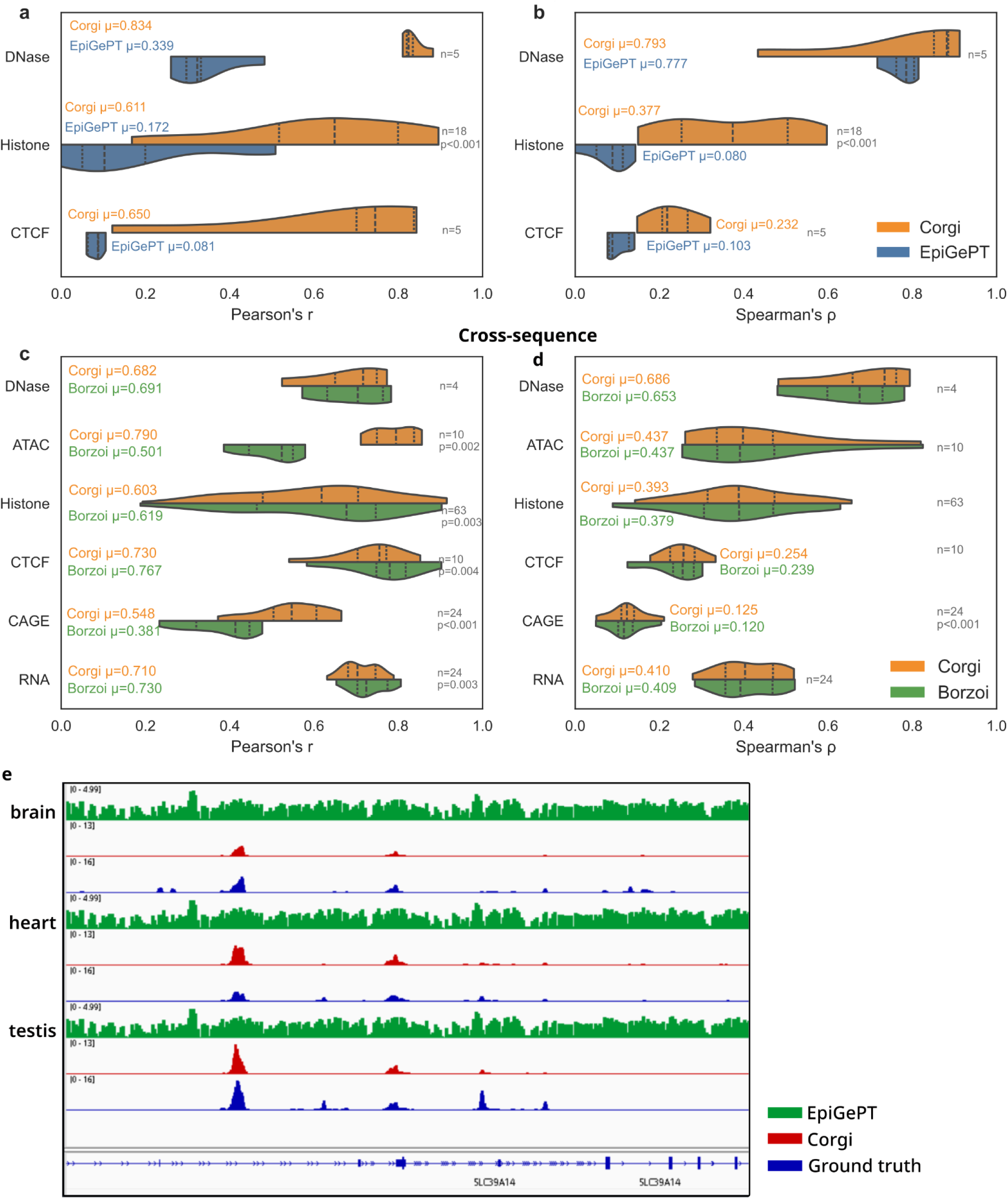
Comparison to EpiGePT and Borzoi. **(a)** Distribution of Pearson and **(b)** Spearman correlation coefficients comparing Corgi and EpiGePT. Sample size corresponds to different samples, and statistically significant results (Wilcoxon signed-rank test) were reported at p < 0.05. Average results are reported next to the violin plots. **(c) and (d)** Similar plots comparing Corgi and Borzoi in a cross-sequence setting. **(e)** Example locus showing EpiGePT (green) and Corgi (red) predictions versus the ground truth DNase-seq signal from ENCODE (blue).

We next compared Corgi to the DNA sequence-to-function model Borzoi^3^ in a cross-sequence setting. Overall, performance across the set of tracks was similar (Figures 3c, 3d, S7). Given that discrepancies in Pearson’s *r* for some tracks may reflect minor differences in output signal scaling (squashed scale) between Corgi and Borzoi, we consider Spearman’s *ρ* to be a more robust metric. Nevertheless, we report both metrics for reference. Gene-level expression correlations using Spearman’s *ρ* were 0.86 for both Corgi and Borzoi, reaffirming that cross-sequence performance is comparable (Figure S6 a,b,e). Gene expression predictions show a very high concordance between Corgi and Borzoi predictions, and the two methods show the same pattern of a subset of genes with close to zero measured expression and nonzero predictions (Figure S6 c,d).

### Corgi can impute epigenomic tracks in a cross-cell type setting

We assessed the imputation performance of Corgi by comparing its accuracy to Avocado, a deep tensor factorization method that can also impute missing data. Unlike Corgi, Avocado does not explicitly use DNA sequence as an input, but instead learns embeddings of genomic positions at three resolutions. We trained Avocado on the ENCODE training set, with RNA-seq tracks from the validation and test sets added in. In this way, Corgi and Avocado can be compared in the setting where both use RNA-seq to predict epigenomic tracks. Corgi has access to only *trans*-regulator expression values, while Avocado has access to the RNA-seq coverage over the training regions.

Corgi shows increased performance over Avocado, with a mean Spearman coefficient of 0.424 compared to 0.362 (Figure 4a). Corgi has a higher prediction accuracy on 63% of the tested experiments, with the largest difference coming from RAMPAGE experiments.

**Figure 4.**
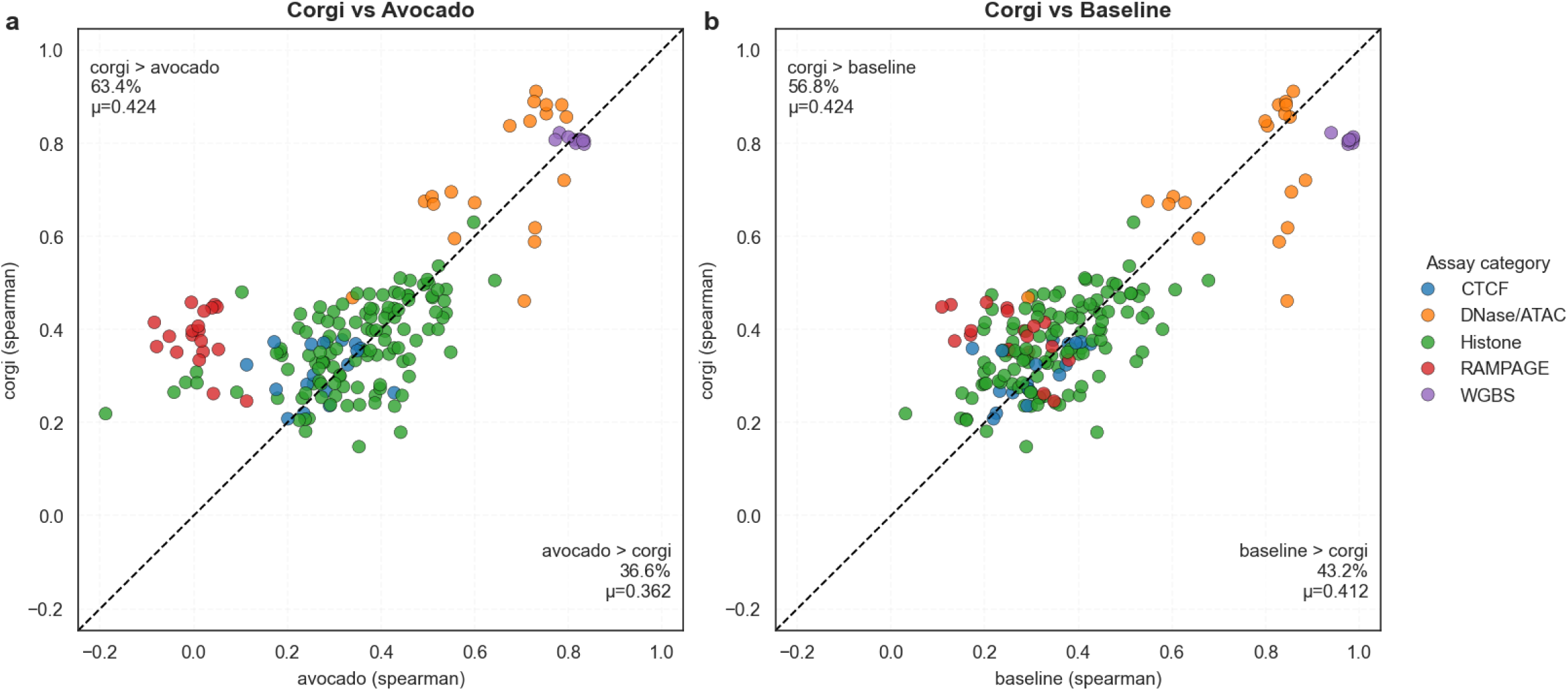
Data imputation performance. **(a)** Scatterplot comparing imputation performance of Corgi vs Avocado and **(b)** Corgi vs Baseline. Spearman correlation coefficients are reported. Each point represents one assay in the cross-cell type set, colored according to the assay category. The dashed line represents the points where performance is equal and it divides the figures in two sections. Mean performance and percentage of points falling into each section is reported.

Given that data from training contexts is available in the same genomic regions in the cross-cell type setting, a baseline prediction can be made by taking the mean signal of all training samples for each assay. Comparing Corgi to this baseline, we see that Corgi has a slightly higher average performance (0.42 vs 0.41) and a higher accuracy on 57% of tested experiments (Figure 4b).

### Corgi understands cell type-specific regulation by learning biologically meaningful *trans*-regulators

So far we have focused on Corgi’s predictive performance alone and have shown that it can make accurate predictions in held-out contexts and sequences. However, genomic sequencing data is highly correlated across different contexts, and often simple baselines can yield good prediction accuracies.

In order to measure Corgi’s ability to uncover cell type-specific regulation, we calculated mean-subtracted gene-level correlations, where the task is to predict the difference of gene expression from the mean gene expression across contexts. In this challenging task, Corgi has a mean Pearson’s r of 0.38 in the cross-cell setting, 0.40 in the cross-sequence setting, and 0.29 in the cross-both setting, showing that it is able to partly understand cell type-specific regulation (Figure 5a). For example, in the embryonic skin tissue from the cross-both benchmark, we can see a high concordance between predicted and measured levels of gene expression differences (Figure 5b).

**Figure 5.**
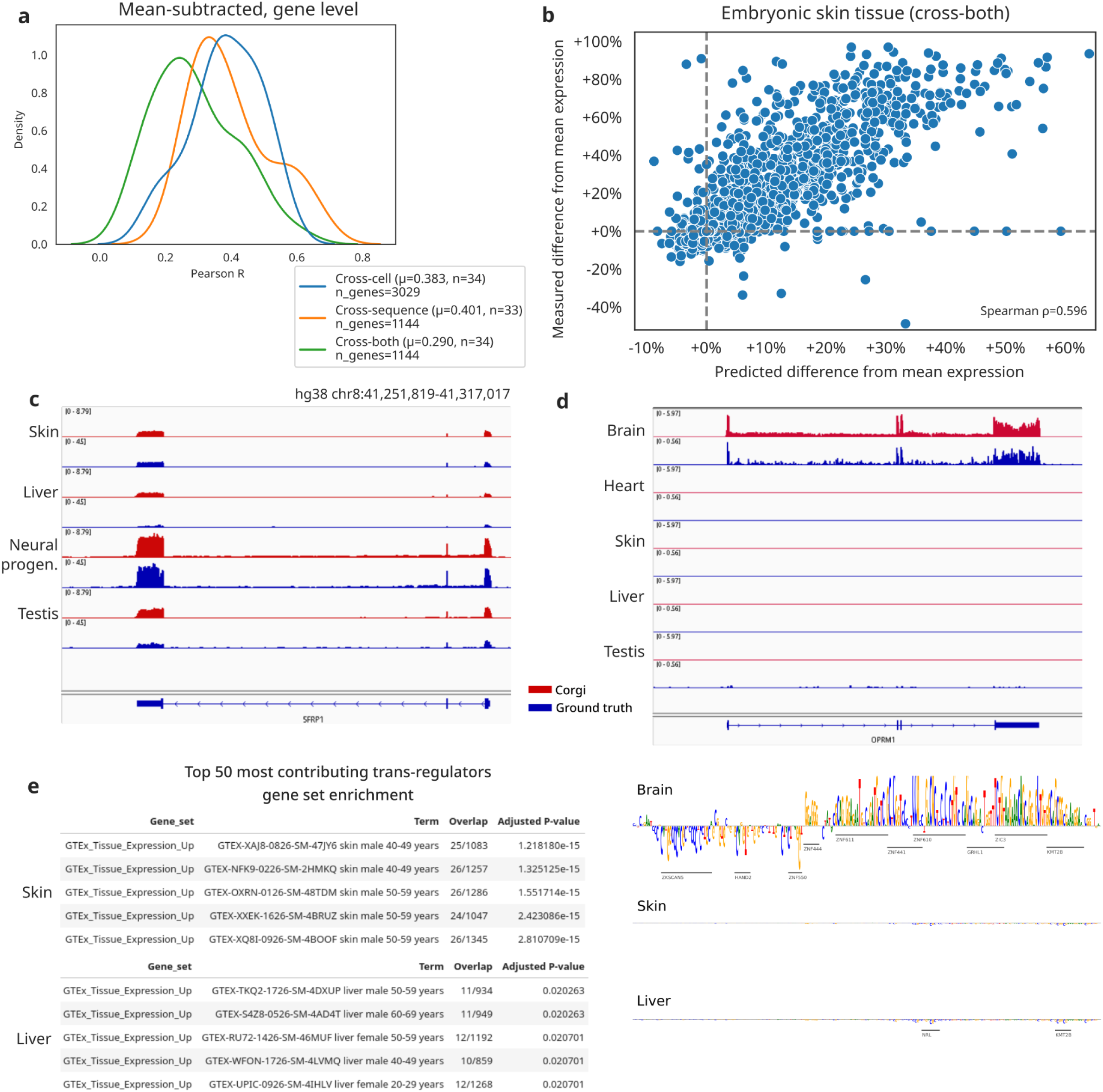
Corgi can understand cell type-specific gene regulation. **(a)** Density plot showing Pearson’s R values of mean-subtracted gene-level correlations under the three testing scenarios. **(b)** Scatterplot showing predicted vs measured difference from mean expression in the embryonic skin tissue. **(c)** Genome browser view around the gene SFRP1, showing predicted (red) and measured (blue) RNA-seq coverage from different tissues and cell types. Corgi correctly identifies neural progenitor cells as the highest-expressed context. **(d)** Genome browser view around the gene OPRM1, showing predicted and measured RNA-seq coverage. Nucleotide contribution scores and annotated TF binding sites around the transcription start site are shown from three different tissues. **(e)** Gene set enrichment analysis results from top 50 most-contributing *trans*-regulators are shown from skin and liver. Corgi identifies upregulated *trans-*regulators in a zero-shot manner.

For concrete examples around specific loci, we looked at RNA expression at the held-out SFRP1 locus, a regulator of neural development in the mouse midbrain^24^, in skin, embryonic liver tissue, neural progenitor cells and adult testis tissue, all of which are held-out contexts (Figure 5c). Corgi correctly predicts gene expression levels of SFRP1 and its upregulation in neural progenitor cells. At the OPRM1 locus, Corgi correctly identifies brain tissue as the only context that expresses OPRM1. An analysis of cis-contribution scores shows increased nucleotide contribution scores around the transcription start site in the brain tissue, while nucleotide contribution scores are lower in skin and liver tissues (Figure 5d). In this genomic window, 7 activating TF binding sites are predicted from the nucleotide contribution scores, and 5 out of the 7 TFs are linked with neural development or neurodevelopmental diseases such as deafness and intellectual disability (ZNF611^25^, ZNF441, GRHL1^26^, ZIC3^27^, KMT2B^28^).

To assess whether Corgi has learned meaningful features that contribute to cell type-specific gene expression, we also computed *trans*-contribution scores of RNA-seq predictions for six contexts across test regions. Gene set enrichment analysis using the top 50 most-contributing *trans*-regulators show that top *trans-*contributors are upregulated in their corresponding tissues (Figure 5e).

In this way Corgi can identify upregulated *trans*-regulators in a zero-shot manner in held-out tissues. Further, in order to test whether Corgi has the ability to causally link *cis* and *trans* effects, we scanned genomic regions with known binding motifs of top *trans-*contributors, and compared these motifs to identified seqlets from *cis-*contribution scores. Top contributors did not show an enrichment compared to a random baseline (Figure S8).

## Discussion

In this work, we present a novel approach that has the potential to shift the paradigm in sequence-to-function models from using sequence alone to context-aware models that can generalize to new cell types as well as new DNA sequences. Compared to previous context-aware models, Corgi shows markedly increased accuracy, a larger context window, increased resolution and the most extensive output experiment types.

When compared to sequence-to-function models that work on DNA, such as Borzoi^3^ and AlphaGenome^4^, the context-aware framework offers possible advantages in using the same data in a more organized manner. The multi-task learning framework used by previous tools treats every genomics experiment as an independent observation. However, genomics data is highly correlated across samples: there are constitutively active regions that show a signal in many cell types and different assay types. On the other hand, if the aim is only to predict variant effects, the multi-task framework is computationally more efficient since it calculates all predictions in one forward pass. Note that having separate output channels for each genomics experiment confers some technical advantages as well: batch effects across samples can be absorbed into the weights of the final convolutional layer. In addition, while Corgi’s training framework requires each output assay type to have a reasonable number of samples in the training data, there is no such constraint in the previous framework. This allows non-context-aware models to be trained on a much larger dataset, including mouse sequences and data, and TF ChIP-seq experiments. Due to slight differences in trans-regulator genes, Corgi does not natively support predictions in the mouse genome, however, this could be circumvented by using orthologous genes or by fine-tuning Corgi on the mouse genome. Here we have shown that despite these technical challenges, Corgi matched the performance of Borzoi while having the additional context-aware prediction capability.

Corgi uses cellular context in the form of *trans*-regulator expression, which includes transcription factors, transcriptional co-activators, chromatin modifiers and RNA binding proteins. This set of genes does not represent all *trans-*regulatory elements in the human genome. We purposefully excluded microRNAs and long non-coding RNAs, as including them would limit us to using samples with available total RNA-seq experiments and discard samples with polyA tail-enriched assays, which includes most single-cell protocols.

We believe that the expression of *trans-*regulators is the most “natural” representation of the cell state for sequence-to-function models, as these regulators directly interpret the regulatory genome, modulate chromatin states and also regulate stability of RNA molecules. Other methods have used only transcription factors, ATAC-seq signal or abstract representations of the entire transcriptome, each having certain drawbacks. Using ATAC-seq is a mathematically convenient approach, because it allows the context vector to be simply concatenated to DNA sequence, yielding five channels (four for the nucleotides, one for ATAC). This is a very rich source of information. Such a model could use promoter accessibility as a baseline for gene expression predictions. However, DNA accessibility cannot be easily manipulated experimentally, and public data availability is lower than RNA-seq. Lastly, some models use an abstract representation of the cell state (scooby^14^, DragoNNFruit^29^) which is not directly manipulatable and hard to interpret.

Corgi can use RNA-seq data to impute missing experiments such as ATAC-seq, histone modifications, CTCF binding and DNA methylation. This can be very useful in situations where only RNA-seq data is available in a given context (rare cell type, patient sample etc.) but predictions on other genomic tracks is desirable. Corgi is more accurate than other tested methods or the mean signal baseline in this task. In addition, this allows nucleotide and *trans-*regulator contribution scores to be calculated, which is useful to uncover why a certain imputation is made.

One of the key advantages of using *trans-*regulator expression as the cell state vector is that *trans-*regulators can be experimentally manipulated with relative ease, using plasmid constructs for overexpression, knockdown or CRISPR knockout of key regulators. Therefore, Corgi theoretically has the ability to simulate these perturbation experiments *in silico,* by running the model with a baseline cell type *trans-*regulator expression and manually changing the expression of selected genes to the desired value. However, we were not able to achieve accurate zero-shot perturbation predictions in practice. Corgi predictions are not sensitive to single *trans-*perturbations, with any effects on the output being miniscule. Several explanations for this phenomenon are conceivable. First, the cell state embedding has around 2900 dimensions and a slight change in one of the dimensions is diluted by the high dimensionality. Second, modeling of the perturbations in this manner does not take into account indirect effects of the perturbation on other *trans-*regulators. In some sense, this is akin to reading out the perturbed state only a very short time after the perturbation is administered, without waiting for the cell to reach a new stable state. However, computationally simulating second- or higher-order effects may not be feasible. This lack of generalization to perturbed contexts is not surprising, as *trans-*perturbation effects are out-of-distribution examples which are notoriously hard to predict, with even the most recent deep learning approaches trained on Perturb-seq^30^ data falling behind linear baselines^31^.

Ultimately linked to accurate prediction of *trans-*perturbation effects is learning of causal relationships between *trans*-regulators and DNA sequence. Causal models are increasing in popularity and will be important for *in-silico* screening of genetic and drug perturbations in new contexts^32^. Here, we showed a novel way of building a context-aware sequence-to-function model by integrating DNA sequence and *trans-*regulator expression information. The next step in the evolution of gene regulatory models will likely be the “causal sequence-to-function model”, which should be able to learn the causal links between abundance of *trans-*regulators, their binding to DNA and ultimately gene expression. Important data sources for this could be TF ChIP-seq and Perturb-seq^30^ data, both of which were not used in training Corgi. Building efficient and biology-inspired deep learning architectures to integrate this data in a causal framework will be key.

## Methods

### Data sources

We scanned the ENCODE database for human samples which have any RNA-seq experiment, plus either one of i) CAGE or RAMPAGE data, ii) DNase-seq and at least 1 histone modification ChIP-seq data, or iii) any combination of at least 4 experiments. It is important that all the experiments are performed on replicates of the same sample. Data that did not comply with ENCODE data quality standards and were marked with an error tag were excluded. Six CTCF ChIP-seq samples with very poor data quality per fraction of reads in peaks (FRiP) score were excluded. In total, we identified 373 samples with these constraints (Supplementary Table 5).

We enriched our dataset with CAGE-seq data from the FANTOM5 consortium. In order to extract maximum information from the available data, we manually curated a list of samples where 65 FANTOM5 CAGE-seq experiments were matched with previously selected ENCODE samples. In addition to these, we selected 119 new samples from FANTOM5 and added them to our set (Supplementary Table 6).

In order to capture variation in *trans-*regulator expression more extensively, we expanded our set using single-cell sequencing experiments. We included 71 pseudobulked single-cell RNA-seq samples from Tabula Sapiens that are matched with single-nucleus ATAC-seq experiments from CATlas. ATAC-seq data was downloaded from CATlas directly as bigwig files, and corresponding matched gene expression data from Fu. et al^12^ was used. Finally, we used 10x multiome single-nucleus joint RNA-seq and ATAC-seq data from brain and peripheral blood mononuclear cells. Data was downloaded from 10x Genomics and after processing and generating pseudobulks, 17 samples were added to the set.

In total, we have 580 biological samples in our dataset, representing a wide range of human cells and tissues. The full list can be found in Supplementary Table 1.

### Defining *trans*-regulatory factors

To determine the set of *trans*-regulatory factors, we need to consider all genes that may play a role in gene regulation. More precisely, we need to find the genes that directly or indirectly regulate DNA accessibility, histone modifications, chromosome folding, DNA methylation or RNA expression. We defined this set to be transcription factors, transcriptional co-activators, chromatin modifiers and RNA binding proteins. Transcription factors directly bind to DNA to regulate transcription and thus constitute the first step of interpretation of regulatory regions on the genome. We used the list of TFs as compiled by the Aerts Lab^33^ which includes 1892 TF genes in the human reference genome hg38^34^. Secondly, transcriptional co-activators interact with transcription factors and regulate transcription and chromatin state, without directly binding to the DNA. The list of transcriptional co-activators was curated from the literature and contains 324 genes, including CREB binding protein, genes involved in the mediator complex and TRIM family proteins^35,36^. Third, chromatin modifiers regulate chromatin state via histone modifications and DNA methylation, thus directly influencing many of our output tracks. We used the dbEM database which contains 167 genes, such as histone acetyltransferases and DNA methyltransferases^37^. Finally, we decided to include all known RNA binding proteins into the set of *trans-*regulators. Since these proteins regulate splicing, transport and decay of RNA molecules, they directly influence RNA-seq coverage. We used a manually curated version from the RBPWorld database^38^ which includes 741 genes. After filtering out duplicate entries and genes not contained in the ENCODE RNA-seq processing pipeline, we are left with a set of 2891 *trans-*regulatory factor genes. The full list of genes can be found in Supplementary Table 2.

We have chosen gene expression as a proxy for the in vivo activity of these proteins, due to the high availability of gene expression data. The unit of gene expression is defined as the logarithm of transcript per million (TPM) values, which is ordinarily calculated from RNA-seq experiments.

### Training-test split

In order to prevent any data leakage across training and test sets, we carefully split our samples into subsets that are as dissimilar as possible. First, we organized the 580 samples into 40 clusters through agglomerative clustering according to their gene expression levels, and left out entire clusters out of the training set until a suitable number of validation and test samples were reached (Figure S2). Second, for all selected validation and test samples, similar cell types or tissues still left in the training set were determined by a keyword match and matching samples were taken out of the training set. Lastly, a random sample was selected from all training clusters into an “easy-test” set, for a testing scenario where the exact samples are unseen by the model, but similar cell types and tissues were seen during training.

In summary, we have 488 training samples, 28 validation samples, 27 easy-test samples and 37 test samples. The test set mainly consists of liver tissue, hepatocytes, testis, prostate, skin, keratinocytes and embryonic stem cell lines.

The genomic sequences were split according to Borzoi’s folds^3^, with fold 3 reserved for testing and fold 4 reserved for validation similar to Borzoi. Furthermore, regions overlapping with gene bodies of *trans-*regulatory factor genes were excluded from the validation and test folds, as their TPM values are used in the input. This resulted in 11,496 sequences of length 524,288 bp for training, 1462 sequences for validation and 1437 for testing, in a tiled fashion across most of the human reference genome (Supplementary Table 3). In total, we train on 2.2 Gbp of DNA sequence and test on 292 Mbp.

### Data preprocessing

The model uses expression levels of *trans-*regulators in units of log(TPM). Importantly, the diversity in gene expression experiments in our training data needs to be addressed because of systematic differences in TPM distributions across different assays. We have 4 types of gene expression assays in our training set: total bulk RNA-seq, polyA-capture bulk RNA-seq, pseudobulked single-cell RNA-seq and CAGE-seq. Since we want to represent biological context, or the cell state, as a universal *trans-*regulator expression vector, ideally this should not depend on the type of gene expression assay that was performed.

In order to control for batch effects across different assays, we harmonized gene expression TPMs of coding genes via taking the logarithm, then quantile normalization and finally batch correction using pyComBat^39^. Quantile normalization and batch correction was performed by using the total RNA-seq data as reference, as this is the most abundant type of gene expression assay in our data. We make this reference available, which allows users to directly use TPM measurements from common RNA-seq assays without the need for further preprocessing, and the Corgi pipeline quantile normalizes them to the reference distribution.

Output signals training data: ENCODE, FANTOM5 and CATlas data was downloaded as processed in the bigwig format. For the 10x multiome experiments, data was downloaded in the h5ad and BAM formats. This data was preprocessed using scanpy^40^, celltypist^41^, multiVI^42^, sinto^43^ and deepTools^44^ in order to generate separate bigwig coverage files for each pseudobulk.

In order to normalize coverage signals across experiments with different dynamic ranges, coverage values were scaled and soft clipped with assay-specific parameters similar to the processing pipeline of Borzoi. Afterwards, coverage values were aggregated at a 64 bp resolution by taking the sum of CAGE and RAMPAGE tracks, the square root of the sum for RNA-seq tracks and the mean for the rest of the tracks.

### Model architecture

The transformer block consists of nine layers that all use multi-head attention at a 64 bp resolution, which represents a good balance between memory cost and prediction quality, while avoiding the need for upsampling techniques such as the U-net used in Borzoi. We use FlashAttention v2 which has speed and memory benefits compared to older implementations of multi-head attention. In total, Corgi has approximately 196 million trainable parameters.

Corgi predicts coverage tracks for the central 393 kb of the input window. The predicted assays are DNase-seq, ATAC-seq, ChIP-seq for multiple histone modifications (H3K4me1, H3K4me2, H3K4me3, H3K9ac, H3K9me3, H3K27ac, H3K27me3, H3K36me3, and H3K79me2), CTCF

ChIP-seq, DNA methylation (whole genome bisulfite sequencing) and transcriptomic assays (CAGE-seq, RAMPAGE-seq, total RNA-seq, polyA-enriched RNA-seq, 10x scRNA-seq). Transcriptomics tracks are strand-specific (i.e. reads aligning to the positive and negative strands are split), with the exception of scRNA-seq. In total, Corgi has 22 channels in its final layer.

### Model training

We trained Corgi with an adapted version of the Poisson multinomial loss function similar to Borzoi, which decomposes the loss to magnitude and shape terms and allows for weighting the shape loss. In order to standardize the loss values across different output tracks, we balanced the loss by weighting each channel’s loss with a learnable weight parameter, inspired by Kendall et al^45^. Thus we minimize a new objective (poisson multinomial * weights + log[weights]) which empirically yields improved performance in some tracks, especially CAGE-seq (Figure S4).

The model was trained for 5 epochs on NVIDIA A100 GPUs with a batch size of 2, for a total of ∼200 GPU-days. We used the AdamW optimizer with learning rate 0.0001 and weight decay 0.01 for the multilayer perceptron and 0.001 otherwise. Mixed precision (bfloat16) was used as it is required by FlashAttention. Furthermore, we used grouped query attention with 4 groups and 8 heads with a total of 192 dimensions. Rotary positional encodings were applied to the first 128 dimensions, similar to Flashzoi^46^.

A cosine annealing learning rate scheduler was used which ramps up over 3000 warmup steps and then decays the learning rate. Due to the lack of computational resources, we did not perform any hyperparameter tuning, which means that the model can possibly be tuned for increased performance.

The number of epochs is considerably lower than most other models, since we consider all pairs of training sequence and training cell type as one training sample and thus our training set size becomes around 5.6 million. In one epoch, each DNA sequence is seen approximately 500 times, which can quickly lead to overfitting before the model has time to learn the rules of cell type specific gene regulation. To combat this redundancy, we applied several dynamic augmentation techniques. First, we use the custom hg38 version from Borzoi in which Gnomad SNPs with a higher allele frequency than the hg38 reference allele were edited in. For all such alleles overlapping an input DNA sequence, each one is randomly edited in with a probability of 0.5. The reasoning is that such alleles could have been the reference allele, and they would mostly have small effects on chromatin and gene regulation, if any. Afterwards, as in previous work, input DNA sequences were randomly shifted up to 3 bp to the left or right, and then randomly reverse complemented.

### Benchmarking

We assessed Corgi’s predictive performance by comparing against the ground truth data for each sample in the test set. For binwise correlations in the cross-sequence and cross-both settings, 637 genomic regions of 524 kb were used by tiling the test fold with a stride of 393 kb. For each of these input regions, Corgi predicts genomic tracks with a 64 bp resolution, resulting in prediction values for 8192 genomic bins, which is cropped to the central 6144 bins. Predictions from the 637 genomic regions are concatenated, resulting in 3.9 million values over which Pearson and Spearman correlation coefficients were calculated. For the cross-cell type setting, a random subset of 947 genomic regions were selected from the training folds. Reported correlations are calculated using the squashed scale, similar to the Borzoi analysis pipeline.

We benchmarked Corgi’s performance against three different published tools: EpiGePT^11^, Borzoi^3^ and Avocado^47^. EpiGePT is the state-of-the-art context-aware sequence-to-function model, and it also uses *trans*-regulator expression and DNA sequence. We were unable to run the provided code to train our own version of EpiGePT, and thus we used the online prediction tool at https://health.tsinghua.edu.cn/epigept/. The published model was trained on 104 cell types on hg38, including samples corresponding to tissues from our test set. Additionally, the genomic regions to be trained on were selected randomly (as opposed to leave-one-chromosome-out or homology-controlled folds), which should also give an advantage to EpiGePT. Nevertheless, we compared EpiGePT and Corgi on 5 selected cell types (brain #124, heart #192, skin #213, liver #277, testis #323) on Corgi’s test sequences. Brain and heart samples are from Corgi’s ‘easytest’ set, so Corgi has seen some brain and heart samples during training, but not these exact samples. Skin, liver and testis are from the Corgi test set. All five cell types were included in EpiGePT’s training set, giving EpiGePT a further advantage in this benchmark. Due to limitations of the EpiGePT’s webserver, only 5 cell types could be tested and a subset of the test genomic regions were used, corresponding to 163,304 bins of 128 bp from chr8. Since Corgi has a 64 bp resolution, average pooling was applied to neighboring bins to reach a 128 bp resolution.

To ensure a fair comparison between Corgi and Borzoi in the cross-sequence setting, we selected 30 samples from our training set that matched specific tracks in the Borzoi training set (Supplementary Table 4). Both models were trained on the same sequence folds, and fold 3 was used for testing, as before. Since Borzoi predicts at a 32 bp resolution, we applied average pooling to match the 64 bp resolution used by Corgi. The aggregation of neighboring bins in this fashion should not decrease the performance, according to the results from the original paper (see Figure 1 from Linder et al. 2025)^3^. We used the Borzoi ensemble (4 models) from https://huggingface.co/johahi/borzoi-replicate-0 (0 through 3) and took the mean of the four model replicates as the final prediction.

Avocado was trained on the same subset of genomic regions from the training folds, using the ENCODE subset of the Corgi training data. Since Avocado needs at least one modality to extract cell type features, RNA-seq data from test cell types were included in the Avocado training set as well. Because we use the RNA-seq bigwigs directly here, samples with harmonized *trans-*regulator expression input from non-RNA-seq samples (CATlas and FANTOM5) had to be excluded from Avocado training. Therefore, we excluded CAGE and ATAC samples from this analysis, since the excluded samples exclusively contain these assays. Every 100 epochs, model performance was evaluated on the validation set, and the best model was chosen at 2900 epochs. We also compared the two models against the training baseline, which is the mean signal from training cell types.

### Contribution score calculation

Nucleotide contribution scores (*cis*-contributions) were calculated using the Captum^48^ package. For each context, the *trans*-regulator expression vector was fixed, and the contributions of individual nucleotides with respect to the sum of the total RNA-seq tracks were calculated using the DeepLift^49^ implementation from Captum. As baseline, the sequence was shuffled with dinucleotide frequencies preserved. Mean contribution scores of 20 random shuffles are taken as the final score. *Trans-*regulator contribution scores (*trans-*contributions) were calculated in a similar way, with mean expression across all contexts as the baseline.

*Cis*-contributions were analysed using tangermeme^50^. First, stretches of highly contributing nucleotides (seqlets) were identified using recursive_seqlets, and then the identified seqlets were matched to known TF binding motifs using annotate_seqlets. HOCOMOCO v13 Core was used as the TF binding motif database^51^. Gene set enrichment analysis of top-contributing *trans-*regulators was performed using the enrichr^52^ implementation from GSEApy^53^.

## Data availability

ENCODE and FANTOM data download links can be found in Supplementary Tables 5 and 6, respectively. CATlas data were downloaded in the bigwig format from https://descartes.brotmanbaty.org/bbi/human-chromatin-during-development/. Processed Tabula Sapiens RNA-seq data matched with CATlas cell types were downloaded from s3://2023-get-xf2217/get_demo^12^. 10x multiome data was downloaded from https://www.10xgenomics.com/datasets/pbmc-from-a-healthy-donor-no-cell-sorting-10-k-1-standard-2-0-0 and https://www.10xgenomics.com/datasets/frozen-human-healthy-brain-tissue-3-k-1-standard-2-0-0.

## Code availability

The Corgi source code, including the model, data processing and training scripts can be found on GitHub at https://github.com/ekinda/corgi. Code to reproduce the results and figures from this paper can be found at https://github.com/ekinda/corgi-reproduction.

## Acknowledgements

We thank Ibrahim Ilik for providing an extended list of RNA binding proteins. We thank Jacob Schreiber for helpful discussions regarding benchmarking. We thank the IT department of the Max Planck Institute for Molecular Genetics and the Max Planck Computing and Data Facility for the usage of computing resources. Figure 1 was made with BioRender with an institutional license.

## Author information

### Author contributions

E.D.A. conceived the Corgi method, developed the computational framework and wrote the initial manuscript draft, with feedback and supervision from M.V. E.D.A. and M.V. wrote the final manuscript.

## Ethics declarations

The authors declare no competing interests.

