## Supplemental Figures for "Context-aware sequence-to-function model of human gene regulation"

### Supplementary Figures

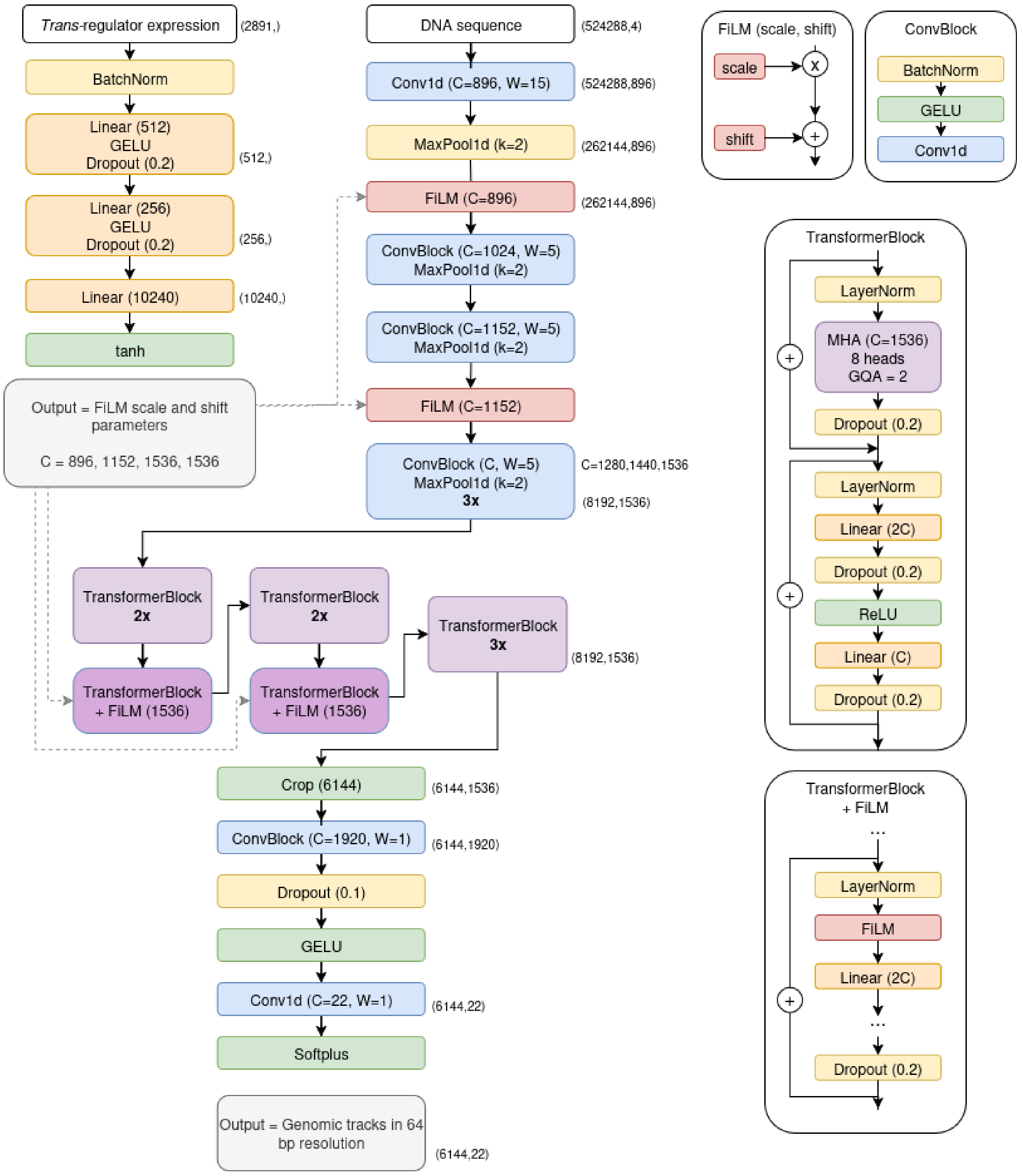


**Figure S1. Corgi's architecture integrates DNA sequence with *trans*-regulator expression**

C = channels, W = convolutional kernel width, k = max pooling kernel width, GQA = grouped query attention. Dashed lines represent information flow to FiLM layers. FiLM layers themselves don't have learnable parameters, they apply an affine transformation to the input based on scale and shift parameters, which are calculated by the multilayer perceptron module.

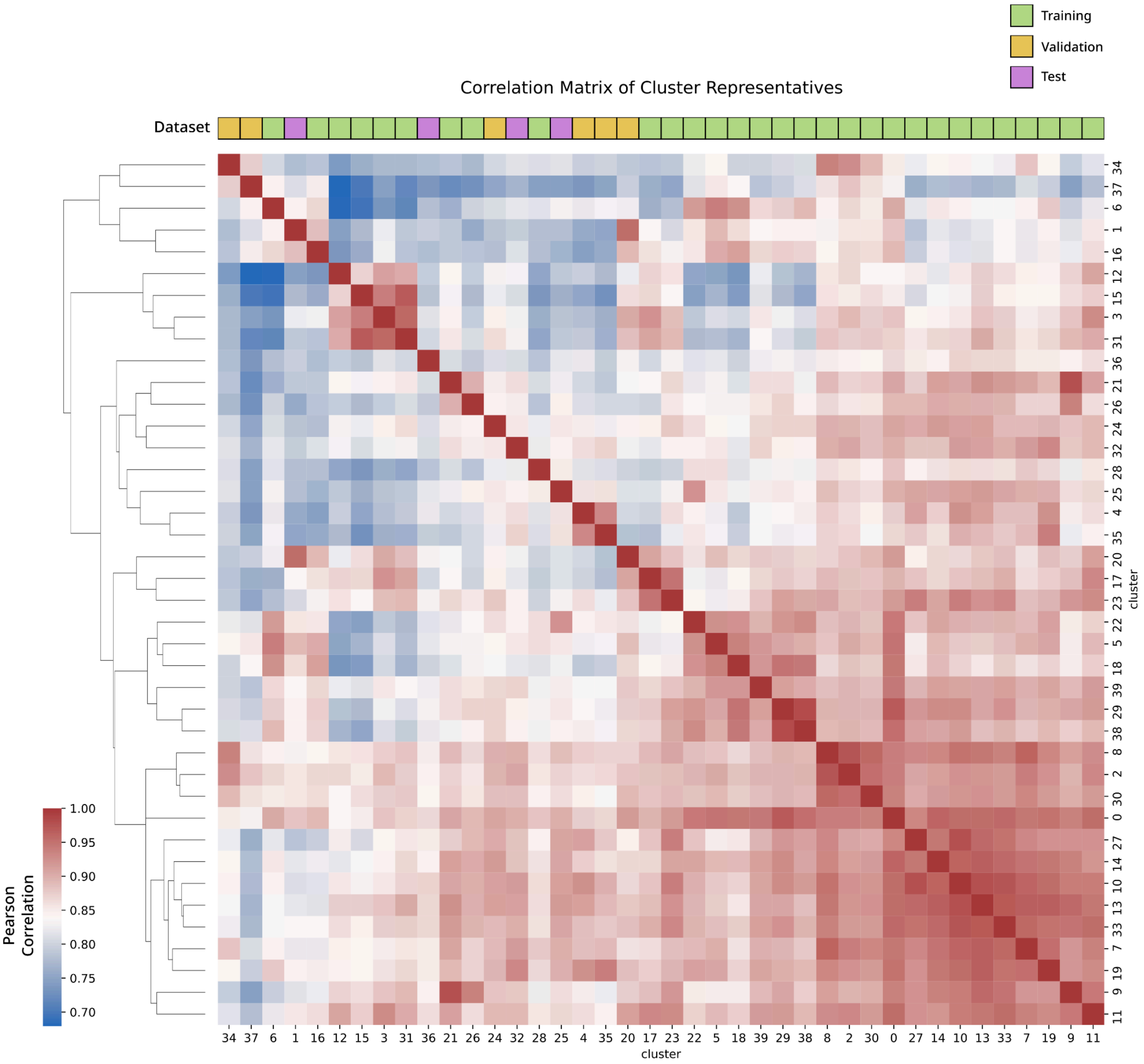


**Figure S2. Similarity matrix of cluster representatives for the training/validation/test split**

The heatmap shows Pearson's correlation coefficients between cluster representatives, after clustering of 580 samples into 40 clusters using an agglomerative clustering based on their gene expression values. Cluster representatives are calculated by taking the mean gene expression of all samples in a cluster. The dendrogram represents hierarchical clustering of cluster representatives, calculated by Euclidean distance. Training, validation and test samples are labeled above the heatmap. We selected validation and test clusters to minimize data leakage and variety of cell and assay types in all three datasets.

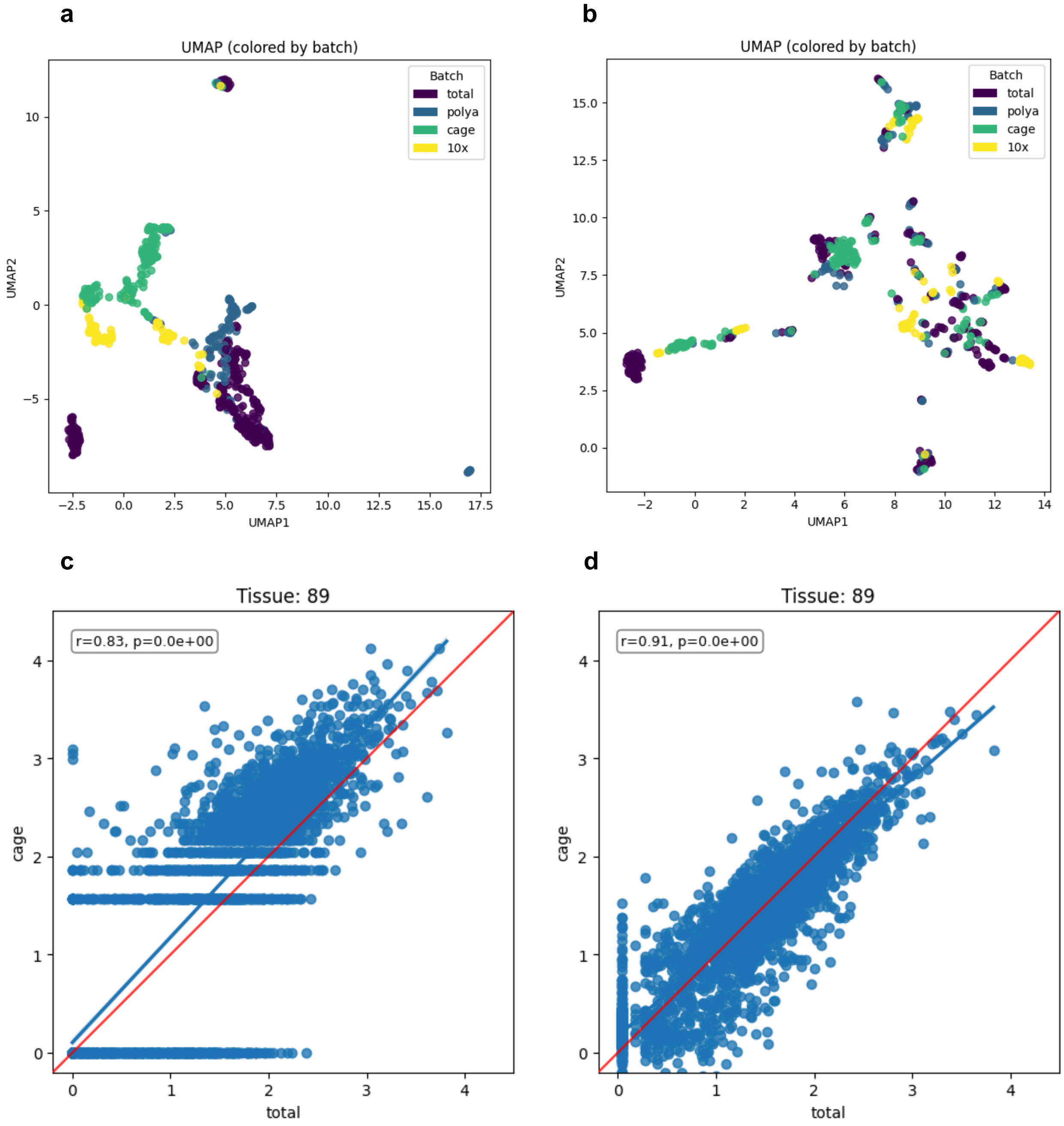


**Figure S3. TPM harmonization reduces batch effects and improves within-sample correlation of gene expression**

(a) and (b) show UMAP plots of gene expression experiments, limited to samples with at least two different assay types available (e.g. RNA-seq and CAGE), for the raw data (a) and harmonized data (b). We see that the raw gene expression values cluster strongly according to their assay types, rather than their cell type. Harmonized values show markedly reduced batch effects. (c) and (d) show the relationship between RNA-seq (x-axis) and CAGE (y-axis) an example sample (#59, K562) which has total RNA-seq and CAGE data available. Raw data (c) has a Pearson's *r* value of 0.83, and the data points do not lie on the x=y line (red). After harmonization (d) we see that correlation improves to 0.91 gene expression calculated by the two assay types are closer to being equal.

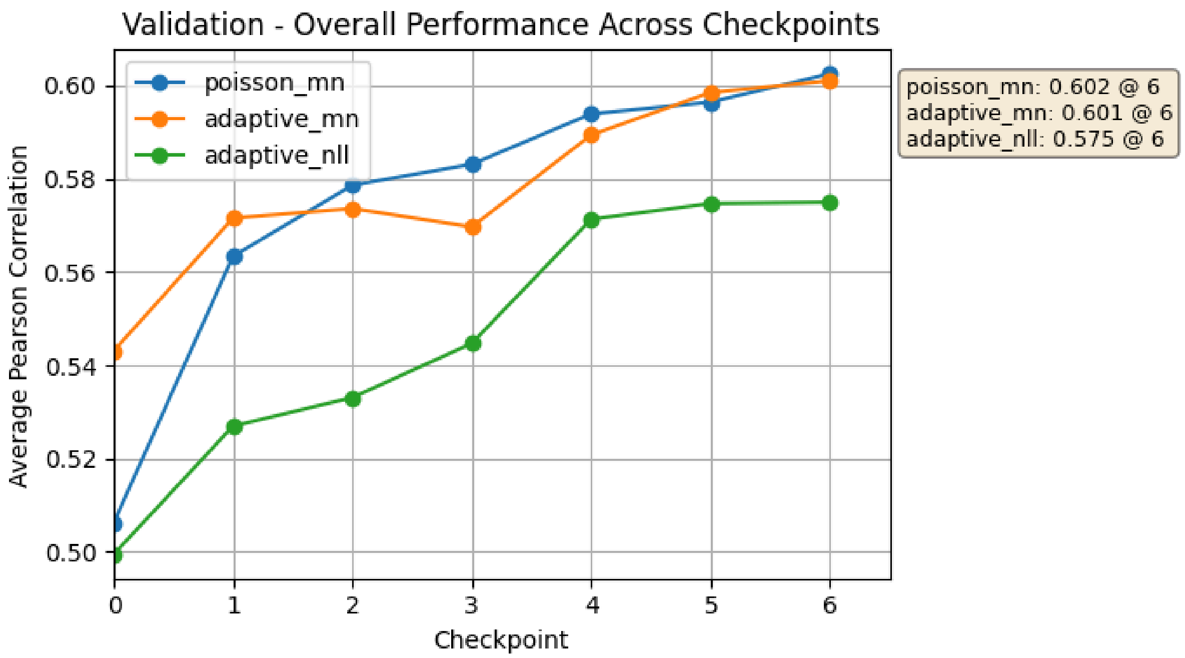


**Figure S4. Validation performance across different model checkpoints**

This plot shows the mean Pearson's *r* of all channels (y-axis) with respect to increased model training. The colors represent different loss function strategies. Orange line is the adaptive poisson multinomial loss function used in Corgi, blue line is the poisson multinomial loss (similar to Borzoi) but with fixed scaling parameters for all output channels (e.g. losses for DNase and RNA are multiplied by 5, while CAGE is multiplied by 100). The fixed values were set empirically. The green line is a poisson negative log likelihood loss function that also has adaptive weights for output channels, but it does not include the decomposition of the loss into shape and total coverage terms. The scaled multinomial loss and the adaptive multinomial loss showed similar performance after the same training epochs. Eventually the latter was selected due to its principled approach as opposed to fixing arbitrary weights empirically. Without using any kind of scaling, performance of certain tracks were diminished (CAGE-seq).

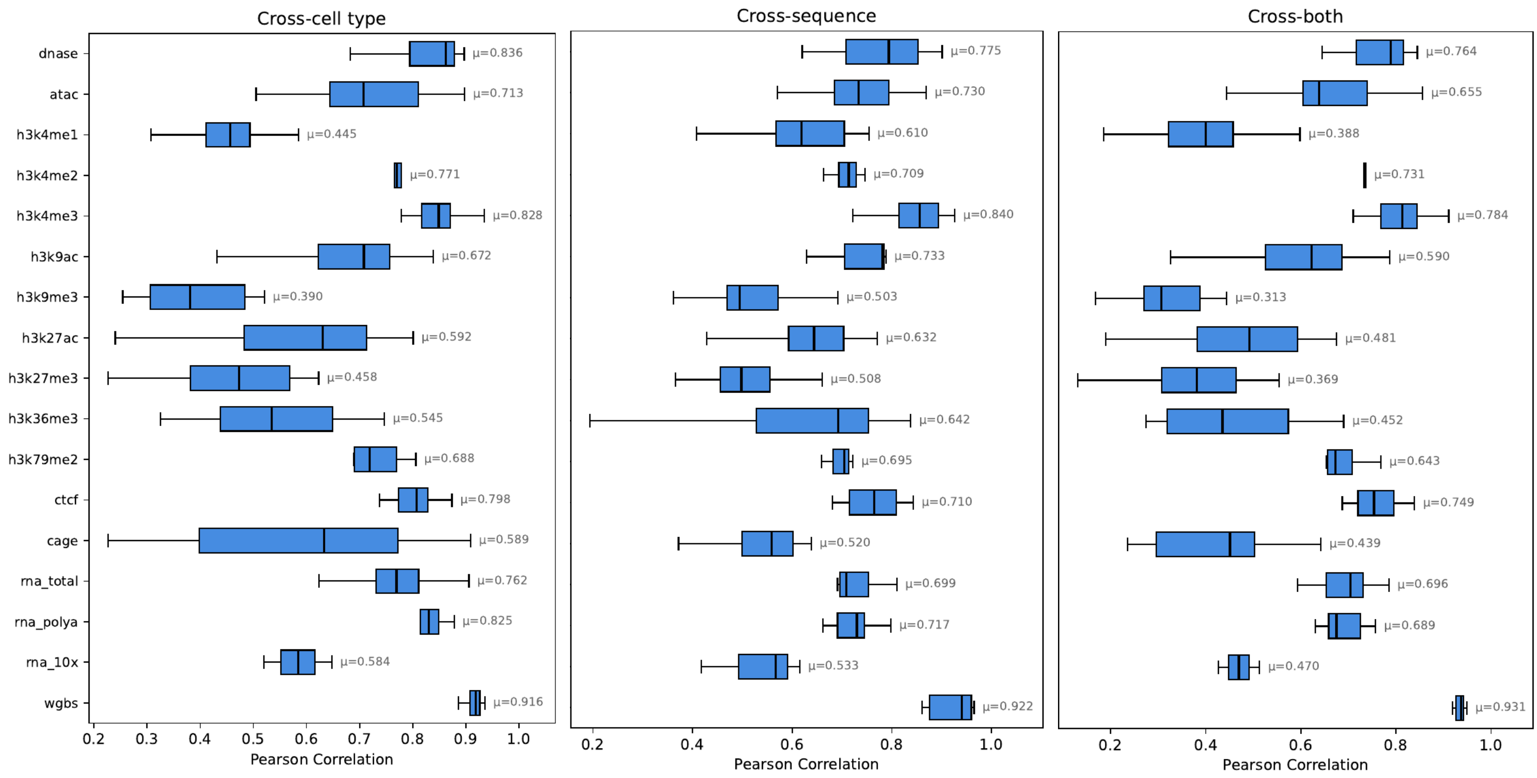


**Figure S5. Detailed genomic coverage prediction performance across different tracks.**

Boxplots showing model performance across different assays in cross-cell type, cross-sequence and cross-both settings. Correlations between predictions and ground truth data across genomic bins are reported, with the variance in the boxplots coming from different biological samples. Pearson's *r* values are reported with the mean values shown next to the boxplots.

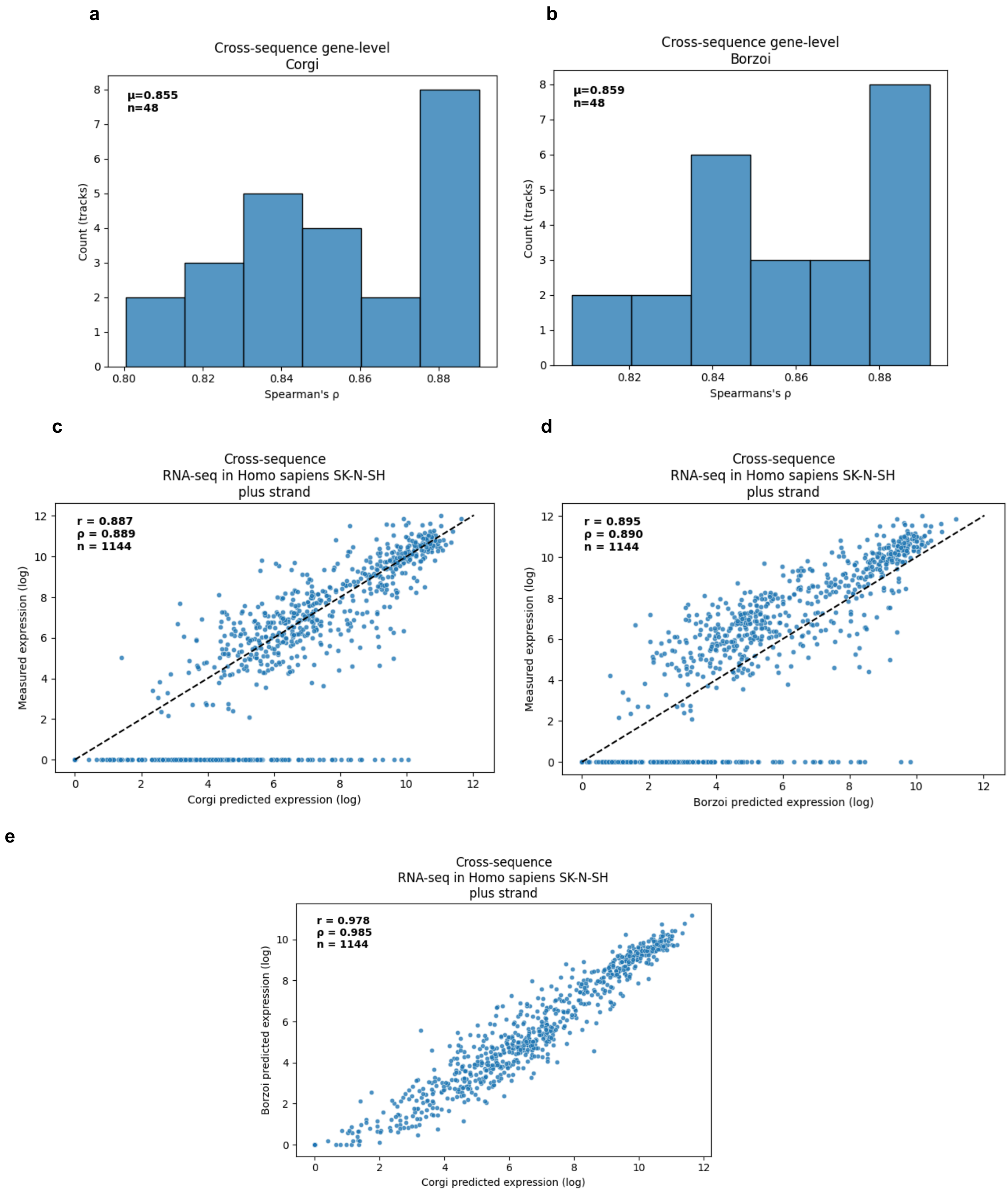


**Figure S6. Gene-level expression prediction performance**

(a) and (b) show distributions of Spearman's ρ across RNA-seq tracks (strand-separated) in a cross-sequence benchmark for Corgi (a) and Borzoi (b). (c) and (d) show an example sample from this benchmark, plus (sense) strand in the cell line SK-N-SH. Corgi (c) and Borzoi (d) perform very well and on a similar level, with both reaching a mean Spearman's ρ value of 0.89. Each point represents a protein-coding gene from the test sequences (Borzoi fold 3). (e) shows that Corgi's and Borzoi's predictions overlap significantly, with a correlation of 0.98. This trend is similar across all samples that were tested in this benchmark.

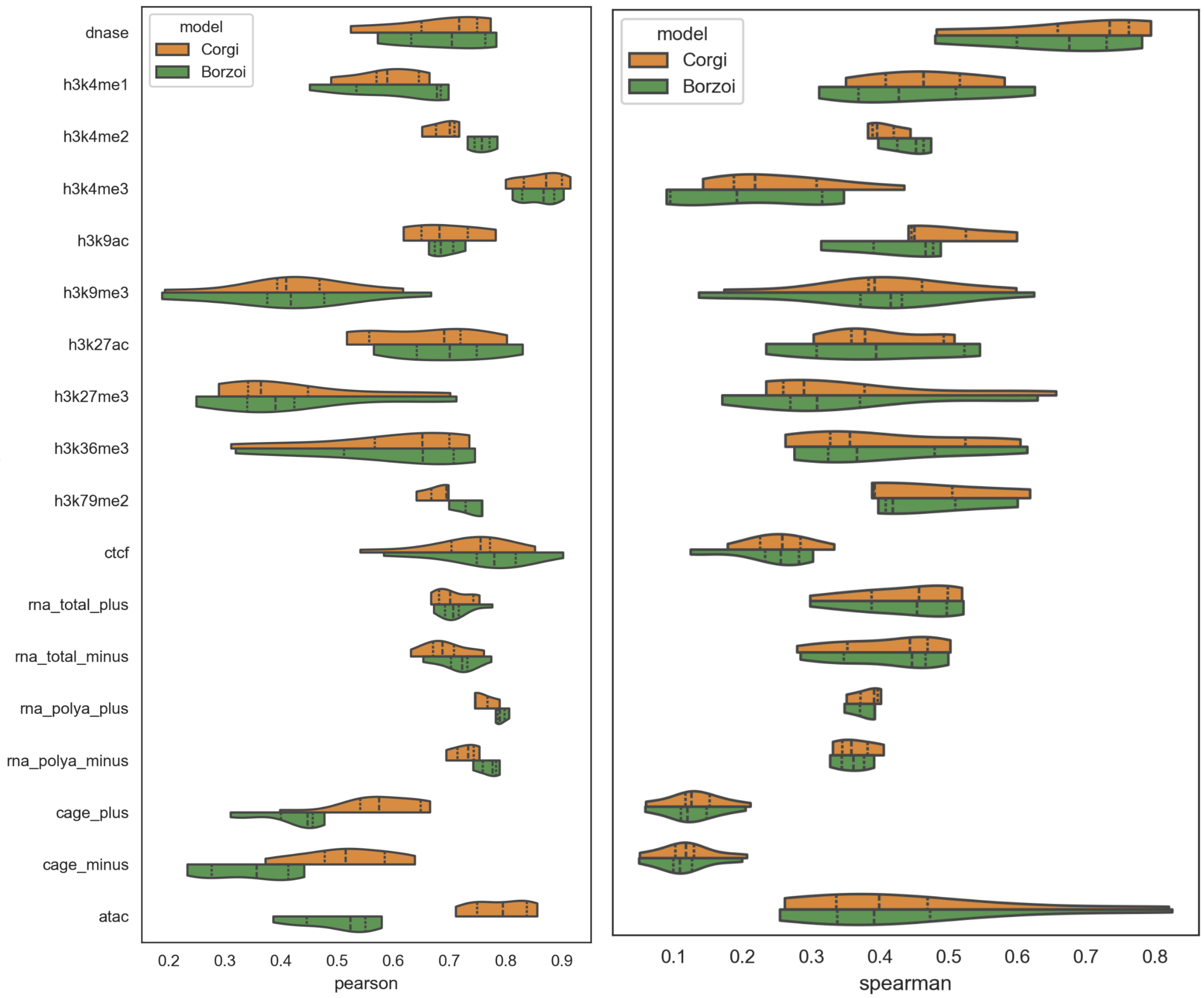


**Figure S7. Detailed performance comparison of Corgi and Borzoi**

Boxplots showing model performance across 30 matched tracks in a cross-sequence setting. Pearson's *r* and Spearman's ρ values are visualized, reflecting correlation between model predictions and ground truth data across genomic bins. Corgi and Borzoi show comparable predictive performance across tracks.

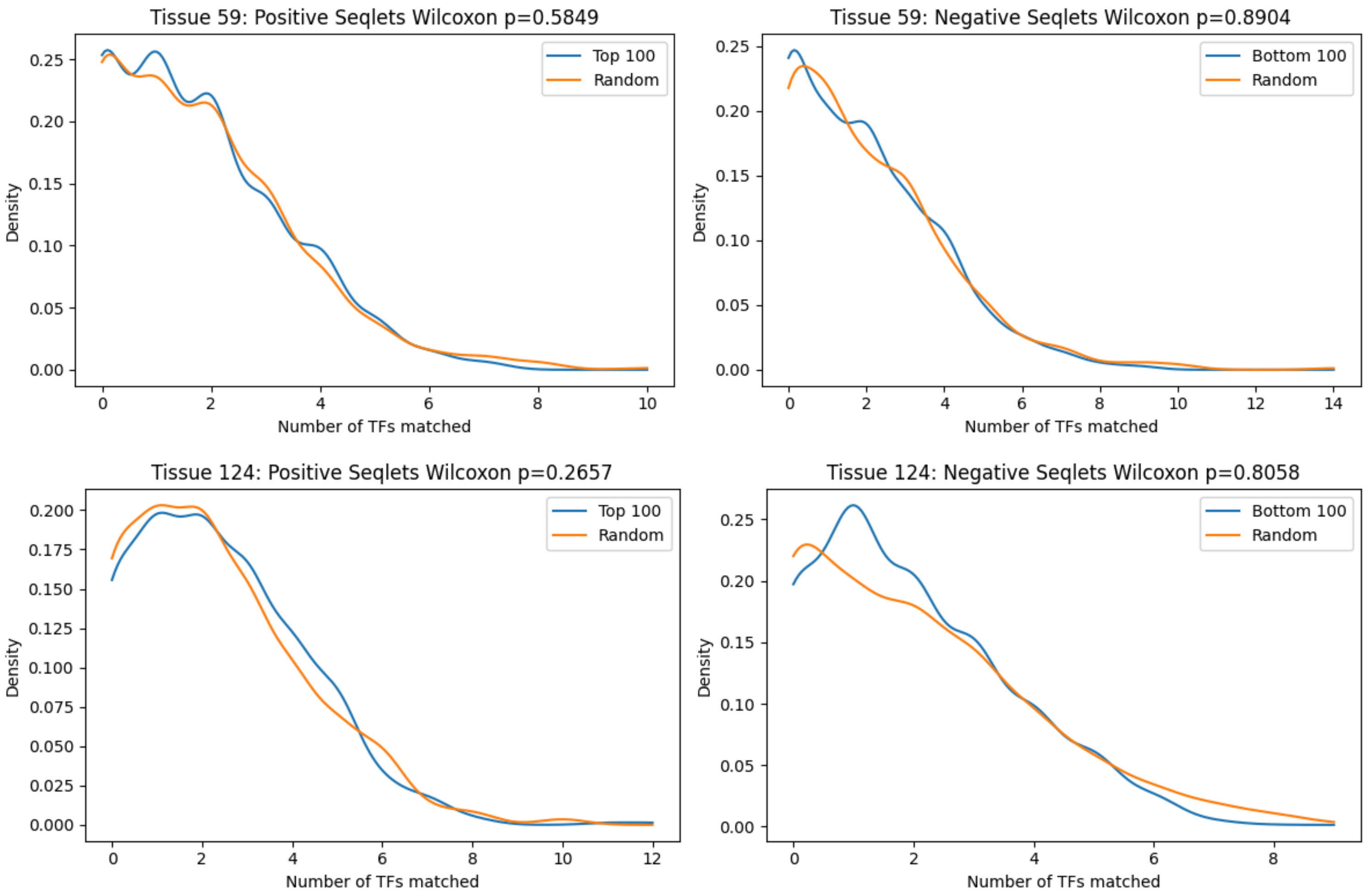


**Figure S8. Matching *cis*- and *trans*-contributors**

Plots comparing the number of transcription factors whose binding motif is highly similar to a seqlet. In the left column, matches from top 100 *trans*-contributing TFs (blue) are compared against a baseline of 100 random TFs. In the right column, bottom 100 *trans*-contributing TFs are compared against the baseline. Top TFs were matched against positively contributing seqlets, while bottom TFs were matched against negatively contributing seqlets. Two example tissues (#59: K562, #124: brain) are shown.
